# Meso-scale multi-material fabrication of a Synthetic ECM Mimic for In vivo-like Peripheral Nerve Regeneration

**DOI:** 10.1101/842906

**Authors:** Paul Wieringa, Ana Rita Gonçalves de Pinho, Roman Truckenmüller, Silvestro Micera, Richard van Wezel, Lorenzo Moroni

## Abstract

A growing focus and continuing challenge for biological sciences is creating representative *in vitro* environments to study and influence cell behavior. Here, we describe the synthetic recreation of the highly ordered extracellular matrix (ECM) of the peripheral nervous system (PNS) in terms of structure and scale, providing a versatile 3D culturing platform that achieves some of the highest *in vitro* neurite growth rates so far reported. By combining electrospinning technology with a unique multi-material processing sequence that harnesses intrinsic material properties, a hydrogel construct is realized that incorporates oriented 6 μm-diameter microchannels decorated with topographical nanofibers. We show that this mimics the native PNS ECM architecture and promotes extensive growth from primary neurons; through controlled variation in design, we show that the open lumens of the microchannels directing rapid axon invasion of the hydrogel while the nanofibers provide essential cues for cell adhesion and topographical guidance. Furthermore, these microstructural and nanofibrillar elements enabled a typically bioinert hydrogel (PEGDA) to achieve similar neurite extension when compared to a biocompatible collagen hydrogel, with PEGDA-based devices approaching neurite growth rates similar to what is observed *in vivo*. Through the accessible fabrication approach developed here, multi-material scaffolds were designed with cell-relevant architectures ranging from meso-to nanoscale and shown to support nerve growth to mimic PNS regeneration, with potential for regenerative medicine and neural engineering applications.

## Introduction

Biological sciences are actively developing various 3D cell niches, in order to better study cell responses in a representative *in vivo*-like environment while under controlled *in vitro* culture conditions.^[1–3]^ This includes the peripheral nervous system (PNS), with a number of *in vitro* platforms developed to accommodate the unique morphology and function of neurons. These unique cells are comprised of a cell body and neurites, specialized cellular projections extending from the cell body that transmit electrical signals to and from tissues. *In vitro* studies often focus on mechanisms of neurite growth as well as signal transduction, in order to better understand the developmental and repair processes involved in neural network formation and to study network function and dysfunction.

The native PNS extracellular matrix (ECM) exhibits a favorably permissive environment for directed neurite growth via a highly ordered 3D architecture, forming an array of oriented microchannels that encase axons from 1 and 20 μm in diameter.^[4,5]^ These ‘endoneurial tubes’ are decorated around their tube perimeters by collagen fibrils that align along the axis of the tubes, resulting in a combination of microstructural guidance and nanotopographical cues that greatly facilitates nerve repair. Because of the presence of this micro-/nanoarchitecture, neural autografts (nerve segments harvested from other regions of the body) remain the clinical “gold standard” for critical nerve defect repair.^[6,7]^ To overcome donor material issues,^[8]^ attempts have been made to develop synthetic alternatives to the autograft^[9–11]^ including a recent nerve guide reported by Bozkurt et *al.* that employs a unidirectional freezing process to form oriented 30-μm-diameter microchannels within a collagen matrix.^[12]^ While in vitro evaluation of this nerve guide reported an estimated growth rate of 145 µm/day, which is far less than the >1 mm/day observed *in vivo*,^[13]^ trials in animals and humans revealed nerve regeneration comparable to the autograft.^[14,15]^ Arguably the best approximation yet of the PNS ECM, the extremely promising nerve regeneration exhibited by this design reinforces the emerging consensus that the 3D microenvironment is crucial for optimal cellular function.

*In vitro* studies of neurite growth on planar substrates have shown how the growth cone, the specialized structure at the leading tip of a neurite, guides nerve growth by responding to local haptotactic and chemotactic cues,^[16]^ such as those found in the native ECM.^[13]^ To guide neurites within an *in vitro* environment, devices such as the Campenot chamber^[17]^ and derivative microfluidic platforms form microchannels on top of 2D substrates through which neurites can grow. Towards a more mimetic culturing environment, oriented nanofiber substrates that approximate the ordered fibrous ECM of the PNS have also been used to guide neurite growth via nanotopographical cues.^[18]^ Meanwhile, the simple embedding of neurons within a hydrogel represents the most accessible 3D growth environment currently available, enabling the first *in vitro* observation of axon myelination by Schwann cells.^[19]^ Insightful studies have further combined oriented fibrils within a hydrogel, highlighting the benefits of incorporating guiding nanotopography within a 3D environment.^[20]^ However, the lack of a permissive microarchitecture within these 3D environments can require neurites to actively remodel the hydrogel, a metabolically active process that results in reduced nerve growth.^[21]^ While the culture environments mentioned are important tools to study nerve growth and function, there remains limited access to culturing platforms that combine the synergistic structural, mechanical and biochemical properties of the ECM.^[22,23]^

Here, we describe a unique 3D multi-material system that strategically assembles hydrogel and nanofiber elements into 3D endoneurial-like microchannel structures, creating a meso-scale culturing platform that we show to support rapid *in vitro* neurite growth of primary sensory neurons. The oriented microchannels are on the scale of endoneurial tubes and do not require neurites to remodel the hydrogel matrix. Similar to the endoneurium, aligned nanofibers are incorporated into the wall of the channels to provide crucial cell adhesive substrates and additional nanotopographical cues. This biomimetic tool offers a promising culturing platform to study PNS regeneration and to optimize strategies for PNS repair or other applications such as regenerative PNS interface design and 3D in vitro neural networks. In general, this accessible fabrication strategy is capable of creating modularly-organized multiscale, multimaterial tissue scaffold towards increasingly representative synthetic ECM mimics.

## Materials and Methods

### Hydrogel Characterization

#### Collagen Crosslinking Dynamics

The crosslinking characterization of collagen was determined with a TA Discovery HR-2 parallel-plate rheometer with Peltier element to control sample temperature. A 4 mg/ml collagen gel was loaded between 40-mm-diameter parallel plates maintained at 5 °C to form a 1-mm-thick pregel solution. Under a constant 1% strain and 1-Hz rotation, the oscillatory moduli were monitored as the temperature was increased to 37 °C, with gelation identified by the crossover point of the storage (G’) and loss (G”) modulus.

#### PEGDA Swelling

To evaluate swelling, triplicate samples of polyethylene diacrylate (PEGDA) hydrogels were immersed in demineralized water for 7 days and the mass of the equilibrium swollen hydrogel (M_swollen_) was weighed. The hydrogel was then dried under vacuum overnight at room temperature (RT), and the mass of the dried hydrogel (M_dry_) was weighed. The swelling ratio (Q) was calculated as follows: 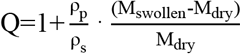 with ρ_p_ the polymer density and ρ_s_ the density of the solvent used to prepare the gel (phosphate-buffered saline (PBS)).

### Device Fabrication

#### Preparation of Polymer Solutions for Electrospinning

A PEOT/PBT block copolymer within the PolyActive™ (PA) family, denoted 300PEOT55PBT45 (300/55/45), was kindly provided by PolyVation B.V. (Groningen, The Netherlands). The chemical composition follows a aPEOTbPBTc notation, where a is the molecular weight, in g mol^−1^, of the starting polyethylene oxide terephthalate (PEOT) segments used in the polymerization process, whilst b and c represent the weight ratio between PEOT and polybutylene terephthalate (PBT) blocks, respectively. PA fibers were fabricated from a 20% w/v solution dissolved in a mixture of chloroform (CHCl_3_) and 1,1,1,3,3,3-hexafluoro-2-propanol (HFIP) at a ratio of 7:3 v/v. Blended fibers were prepared from a PA-collagen solution, which was a 1:1 blend of a 20% w/v PA solution prepared in HFIP and 8% w/v bovine collagen type I (a generous gift from Kensey Nash Corporation (USA, Catalogue number 20003–04)) also in HFIP. This produced a final solution with 10% w/v of PA and 4% w/v of collagen in pure HFIP. The sacrificial template fibers were fabricated from poly(D-L)lactic acid (P(DL)LA) 50/50 (Mw=40000, IsoTis S.A., The Netherlands), dissolved in a mixture of dicloromethane and HFiP (4:1). Concentrations of 50% w/v and 75% w/v were prepared. All solutions were prepared and stirred overnight at RT before use.

#### Preparation of Nano- and Microfiber Templates

Electrospun fiber templates were prepared using a custom apparatus with temperature and humidity control, maintained between 24–25 °C and 28–32% rH, respectively. The polymer solution was loaded into syringes mounted on a syringe pump (KDS 100, KD Scientific). A stainless-steel spinneret (with an outer diameter (OD) of 0.8 mm and inner diameter (ID) of 0.5 mm (Unimed, Lausanne, Switzerland) was mounted to an upper parallel plate and connected to a high-voltage generator (Gamma High Voltage Research Inc., FL, USA). Aligned fibers were achieved by employing a gap electrode grounded target, consisting of two aluminium electrodes 2 mm wide with a gap of approximately 15 mm (Figure 1, inset). A Teflon^®^ mount was used to position a ring-like mesh support frame (15 mm OD, 12 mm ID) across the electrode gap. Fibers were collected onto the support frame, with both the fibers and the frame later embedded within the final hydrogel construct. For PA-collagen fibers, we used a flow rate of 0.2 ml/h at a voltage of 15 kV with an air gap of 15 cm. PLA fibers were electrospun at flow rates of 1, 5, and 10 mL/h for the 50% w/v solution, and 5 ml/h for the 75% solution. Voltages ranged from 15 to 20 kV and an air gap between 20 cm and 25 cm. Fiber-only (FO) templates consisted of PA-collagen nanofibers deposited onto a mesh support for 5 min. Channel-only (CO) templates were created by depositing PLA microfibers for 5 s. Templates with microchannels and nanofibers (CF) were formed from a triple layer of alternating fiber deposition of PA-collagen (2.5 min), PLA (5 s), and PA-collagen (2.5 min), followed by a thermal treatment between 60 °C and 80 °C for 1 h.

**Figure 1.**
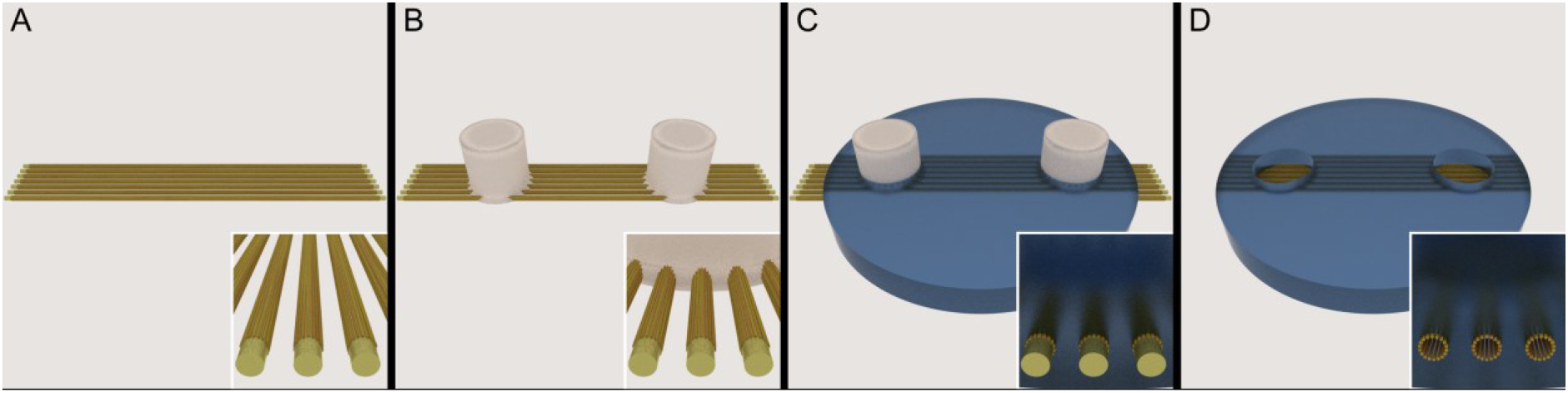
Schematic of the mutlimaterial template fabrication strategy. Using a gap electrode collector, the sequential deposition of aligned fibers resulted in creating a template layer of soluble microfibers (yellow) between two insoluble layers of nanofibers (orange) (A). The multimaterial fiber network was then heated for 1 h at 65 °C, allowing for the insoluble fibers to merge on the periphery of the template fibers (cross-sectional view, A inset). PCL plugs (white) were then deposited onto the fiber network (B), forming the well templates for cell seeding. The fibers were then embedded between layers of hydrogel prescursor (C). After crosslinking of the hydrogel, the construct was place in warmed acetone to dissolve the template, leaving the final hydrogel construct (D) with microchannels lined with aligned nanofibrous topography (D, Inset).

#### PCL Plug Fabrication

Sacrificial plugs were used to form the seeding wells within the *in vitro* hydrogel device. Polycaprolactone (PCL; Capa 6500, Perstorp, UK) plugs were manufactured using a Bioplotter 3D printer (Envisiontec GmbH, Germany). The polymer was melted at 145 °C in a stainless steel syringe with a 1.2-gauge needle, and the molten polymer was extruded via mechanical pressure to merge with fibers. The syringe was raised while additional polymer was extruded, forming a 5-mm high PCL pillar with a 3-mm diameter, creating well templates merged with the underlying aligned fiber matrix (Figure 1B). Two wells per scaffold were created in this manner.

#### Hybrid Hydrogel Device Assembly

Hydrogel scaffolds of either collagen or PEGDA were formed in cylindrical wells of an aluminium mold (17 mm in diameter and 10 mm deep). Starting from a high concentration solution of rat tail collagen in 0.02 N acetic acid (Corning), a 4 mg/ml solution of pre-collagen was prepared in 10× PBS, sterile dH_2_O, and 1 N solution of NaOH and maintained on ice until use according to supplier instructions. A pre-polymer solution of PEGDA (Mw=5000, Laysan Bio) was prepared in PBS (Gibco, Invitrogen) at RT, for a final concentration of 10% w/v. A solution of Irgacure 2959 (BASF) photoinitiator was freshly prepared in 70% EtOH at a stock concentration of 10% w/v and added to the PEGDA at a dilution of 1:100 for a final Irgacure concentration of a 0.1% w/v. 100 µl of pre-polymer solution was dispensed to form a thin hydrogel layer: collagen was placed in an incubator at 37 °C and 5% CO_2_ for 5 min to initiate crosslinking; PEGDA was briefly crosslinked for 2.5 min under N_2_ flow in a crosslinking cabinet (Ultralum, USA) equipped with 365-nm UV lamps (10 mW/cm^2^). Prewetted templates, comprised of a support frame, electrospun fibers, and PCL plugs, were positioned on top of this layer and held down by a tubular stainless steel insert (17 mm OD, 15 mm ID, 2 cm height). To the well, 300 µl of hydrogel was added and crosslinked as before: collagen was allowed to crosslink for 1 h; PEGDA was crosslinked by UV light for 10 min. The template was then dissolved by placing the construct in acetone (Sigma Aldrich) held at 50 °C and mixed gently. The acetone was refreshed every hour for the first 2 h, then maintained for 16 h (overnight). This was followed by the same sequence in 100% ethanol at 37 °C, followed by the same sequence in sterile demineralized water at 37 °C.

### Device Characterization

#### SEM Analysis

Electrospun fibers were gold-coated using a sputter coater (Cressington SputterCoater 108auto) and imaged by scanning electron microscopy (SEM; XL 30 ESEM-FEG, Philips). The diameters of 150 fibers per condition were measured using the ImageJ image analysis software (National Institutes of Health, USA). Hydrated hydrogel devices were cut with a sharp blade and dehydrated in a tert-Butanol (TBA, Sigma Aldrich) dilution series of 50%, 70%, 100%, 100%, 100% TBA/H_2_O for 12 h each, maintained at 37 °C. The samples were frozen in liquid nitrogen and placed under vacuum at 4 °C to sublimate the TBA overnight. The samples were later gold-sputtered and imaged.

#### Microchannel Diffusion Assessment

To verify that open channels were formed within a hydrogel, 20 μl of FluoSpheres^®^ solution (100- and 200-nm diameters, Invitrogen) was added to one well, and diffusion through channels was monitored via time-lapse epifluorescent microscopy (EVOS® FL digital fluorescence microscope). Confocal microscopy (Nikon A1 confocal microscope) of perfused channels was used for 3D reconstruction of microchannels within the gel.

### Primary Cell Culture

Dorsal root ganglia (DRGs) were isolated from postnatal rat pups (Wistar Unilever, HsdCpb:WA) between the ages of 2 and 8 d. All procedures followed national and European laws and guidelines, and were approved by the Netherlands animal ethics committee. Briefly, rats were sacrificed by cervical dislocation under general anaesthesia (4% isoflurane) and then decapitated. Individual ganglia were removed from the spinal column and nerve roots were stripped under aseptic conditions with the aid of a stereomicroscope. Hydrogel devices were incubated in sterile PBS (Invitrogen) for 2 h, followed by placement in a 24-well plate (one device per well), with each well filled with PBS to avoid trapping air bubbles under the hydrogel. The PBS was removed, and hydrogels were held within the well using Viton^®^ O-rings (Eriks BV, The Netherlands). Devices were then incubated in 200 µl of laminin (15 µg/ml; Sigma Aldrich) and poly-l-lysine (0.2 µg/ml; Sigma Aldrich) in PBS for 2 h at 4 °C. Each device was then washed 1× with PBS and 2× with Neuralbasal™-A medium (Invitrogen), with the last wash applied for 1 h. This was replaced with 200 μl of complete Neuralbasal^®^-A medium, supplemented with 0.5 mM of L-glutamine, 1× B27 supplement, and 10 U/ml of penicillin/streptomycin (all from Invitrogen). Medium was additionally augmented with either 50 ng/ml of glial-derived growth factor (GDNF; Sigma Aldrich) or 10 ng/ml of neurotrophic growth factor (NGF; Sigma Aldrich). Each scaffold was seeded with 1 DRG in one of the wells, and cultures were maintained for 5–8 d, with medium changed every 2 d.

### Immunohistochemistry and Imaging

Medium was aspirated from cell culture samples, cells were washed twice with a wash buffer [tris-buffered solution (TBS; Sigma Aldrich) + 1% w/v bovine serum albumin (BSA; Sigma Aldrich) and 3.4% w/v (100 mM, 34 mg/mL) sucrose (Sigma Aldrich)] and fixed for 1 h with a 4% w/v paraformaldehyde solution (Sigma Aldrich) at 4 °C. After 3 times washing with the wash buffer, cells were permeabilized with 0.2% v/v Triton-X 100 (Sigma Aldrich) in wash buffer for 30 min. Next, they were washed again 2 times with the wash buffer and incubated with 5% v/v goat serum (Sigma Aldrich) for 1 h. Primary antibodies *β*-Tubullin III (1:1000, anti-mouse, Sigma Aldrich) and S100 (1:500, anti-rabbit, Sigma Aldrich), both neuron-specific proteins, were diluted in wash buffer with 2% v/v goat serum and incubated for 24 h at 4 °C. Samples were washed 3 times with wash buffer + 2% goat serum, then incubated with goat anti-rabbit IgG conjugated to Alexa 488 (Invitrogen) and goat anti-mouse IgG conjugated to Alexa 594 (Invitrogen) diluted in wash buffer (1:500) for 16 h at RT, in the dark. GDNF-sensitive neurons were stained by first blocking endogenous biotin and avidin (VectorLabs), 30 min for each step, followed by incubation with Isolectin-B_4_(IB4)-biotin (5 µg/ml, Invitrogen) for 24 h at 4 °C in TBS. After a triple TBS wash, samples were incubated for 16 h with streptavidin-488 (Jackson Immunoresearch, USA). To visualize the cell nucleus, cells were incubated for 20 min in 0.7 µg/ml 4′,6-diamidino-2-phenylindole (DAPI; SigmaAldrich) as counterstaining, followed by a 2× wash in wash buffer. Samples were imaged with a confocal microscope (Nikon A1 confocal microscope).

### Statistical Analysis

All statistics and graphs were produced with R statistical software (http://www.R-project.org/). Neurite growth (n≥3) and fiber diameters (n>20) were compared with a one-way ANOVA followed by a *post hoc* Tukey HSD test (p < 0.01). A Student’s T test was used (p < 0.05) to compare template fiber diameter and microchannel diameter. Standard deviations are used unless otherwise stated.

## Results

To realize this hydrogel culturing platform, a combination of a sacrificial microfiber template and insoluble nanofibers was used to form meso- and microstructures that incorporate oriented nanotopographic cues. To facilitate the device assembly, a gap electrode collector was used to deposit aligned electrospun micro- and nanofibers onto a supporting polyester mesh frame that permitted handling of the delicate fibers throughout the fabrication process. Poly-l-lactic acid (PLA) microfibers were optimized to form an oriented microchannel template by altering the electrospinning parameters (flow rates of 1, 5, and 10 ml/h) and the composition of the PLA solution (50% and 75% w/v), resulting in fiber diameters ranging from 2.5 µm and 15.7 μm (Figure 2A); this approximates the diameter range of axons that reside within the endoneural tubes.^[4,5]^ The 50% w/v PLA solution at a flow rate of 5 ml/h achieved a mean fiber diameter of approximately 6.0 μm (Figure 2B) and was subsequently chosen for follow-up experiments to approximate the endoneurial tube diameter of a sensory nerve.^[24]^ Analysis confirmed a well aligned microchannel template based on a coherence measurement of 0.77 (Figure S1), where 0 is completely random and 1 is perfectly aligned.^[25]^

**Figure 2.**
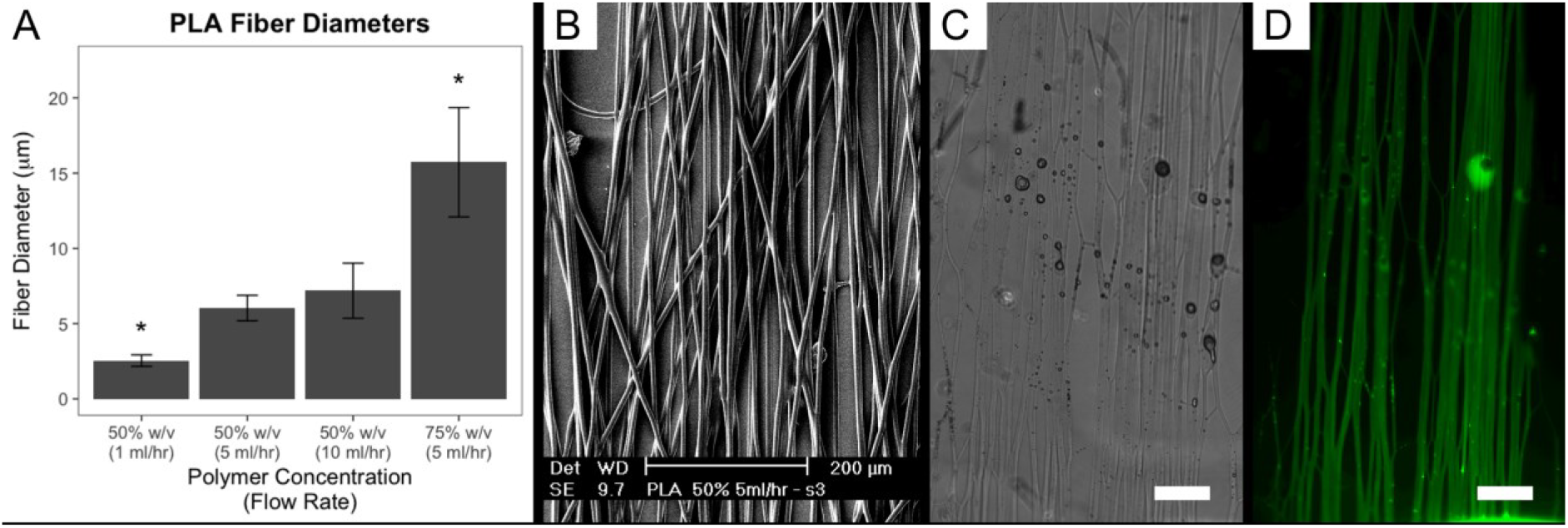
Microchannel characterization. Electrospinning of PLA fibers under different conditions. Three distinct diameter populations were produced (p<0.01; n>20) (A). A SEM image of oriented PLA fibers produced from a 50% w/v solution at a flow rate of 5 ml/hr (B); these microfibers were used further as the microchannel template, with an average diameter of 6.04 ± 0.85 µm. The resulting microchannels could be observed through bright field microscopy (C) and microchannel/well connectivity was confirmed by the diffusion of fluorescent microspheres (D). Also evident is the arborized microchannels that results from the bundling of the template fibers. Scale bars: B, 200 μm; C, D, 250 μm.

Fiber nanotopography was combined with the microchannel template by sequentially depositing aligned fibers to create a triple-layer fibrous construct of PLA microfibers sandwiched between nanofiber layers (Figure 3A). Nanofibers were electrospun from a blended solution of PolyActive™ (PA) and collagen to provide oriented topographical guidance cues. Such a blended composition has previously been shown to improve DRG adhesion^[28]^ and resulted in nanofibers that were sufficiently robust for processing, with an average diameter of 218 ± 57 nm and a coherence of 0.75 ± 0.11 (Figure S2).

**Figure 3.**
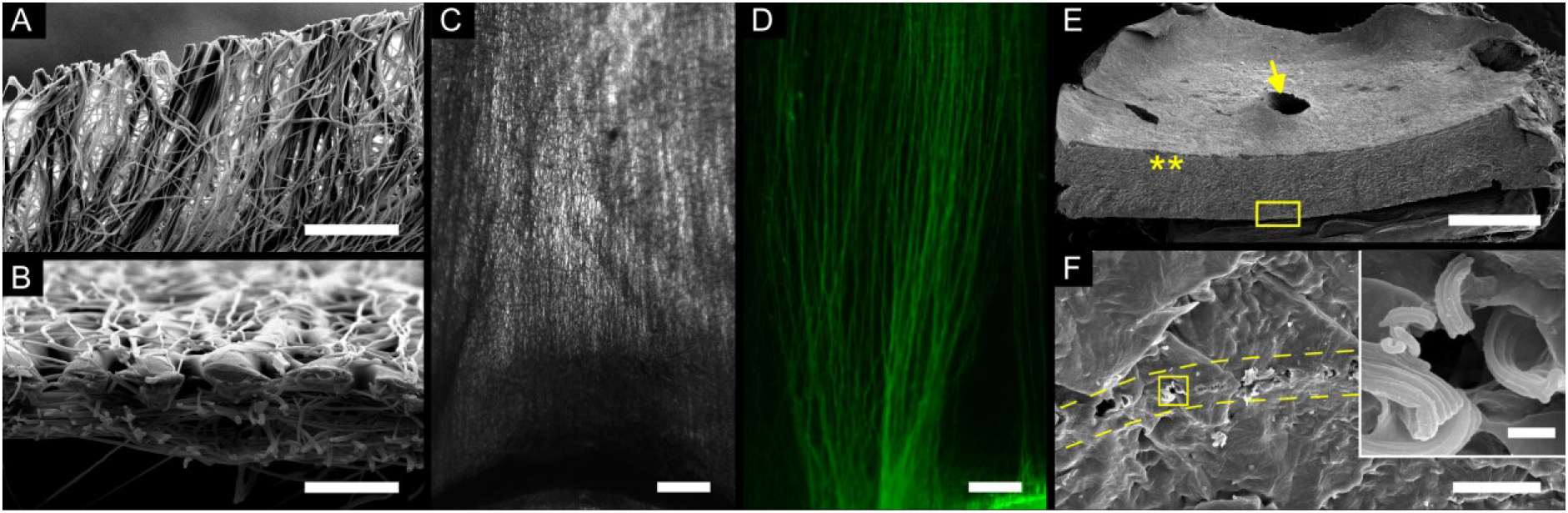
Optimization of the CF template and verification of fiber/channel integration. The triple layer (A) was subjected to a 65 °C thermal treatment, successfully merging the insoluble nanofibers on the periphery of the microfiber template fibers (B). A brightfield image of a PEGDA-CF shows the embedded nanofibers within the hydrogel (C) with fluorescence microscopy confirming the presence of clearly delineated microchannels in the device, demonstrated by the directed diffusion of fluorescent microspheres over a 30 minute period from one mini-well to another (D). SEM images of an assembled PEGDA-CF hydrogel construct that has been bisected (E) shows a device with a mini-well (arrow) and the cross-sectional face of the construct (**). Inspection of the cross-section (E, yellow box) reveals the formation of a row of microchannels (F) located between the yellow dashed lines. Closer examination of the microchannels (F, yellow box) showed they were delineated by nanofibers (F, inset). Scale bars: A 50 μm; B 20 μm; C, D 250 μm; E 1 mm; F 20 μm; F inset 1 μm).

To ensure the nanofiber topography was exposed within the mini-well and inner perimeter of the microchannel wall to allow cell adhesion, a unique thermal and solvent strategy was developed through careful material selection based on solubility, melting (T_m_) and glass transition (T_g_) temperatures. By applying a 65°C thermal treatment to micro-/nanofiber construct for 1h, the PA nanofibers (T_m_: 145°C) could superficially merge at the periphery of the thermally softened PLA microfibers (T_g_: 60 °C) (Figure 3B). This ensured the microchannel template retained a round morphology while leaving the nanofibers exposed within the microchannels after template dissolution (Figure 3F). Following this thermal treatment, fused deposition of polycaprolactone (PCL) formed pillars, 3 mm in diameter and 5 mm high, on top of the fibers to create the mini-well templates to be used for cell seeding (Figure 1B); the relatively low T_m_ of PCL (60 °C) allowed the PLA microfibers (T_m_ = 160 °C) and PA nanofibers to remain intact while being enveloped by the mini-well template (Figure S3). This was critical to form interconnected mini-wells with exposed nanofibers on the bottom surface, such that cells could both adhere to the well and navigate into the oriented microchannels.

The water insolubility of the template structures and guidance nanofibers was critical to ensure both remained intact while being embedded within an aqueous hydrogel precursor. Equally critical was the selective remove of the template structures while leaving both the crosslinked hydrogel and nanofibers intact. Acetone was chosen as the ideal solvent, since it can dissolve both PCL and PLA but not PA, resulting in a highly ordered 3D multimaterial environment with interconnected structural features ranging from the meso-to the nanoscale.

The ability to assemble different components, as described above, makes this a versatile platform to study cell interaction through the systematic variation of structural elements. In the current study, three different types of devices were realized: 1) two wells linked via oriented nanofibers (FO – fibers only); 2) two wells linked via oriented microchannels (CO – channels only); 3) two wells linked by microchannels that incorporate oriented nanofibers around the microchannel periphery (FC – fibers and channels). Furthermore, PEGDA was selected as a representative synthetic bioinert material to highlight the role of incorporating cell-adhesive nanofibers. For comparison, collagen hydrogel was chosen as a representative cell-compatible natural gel, for a total of six different platforms.

PEGDA hydrogel devices were formed in a cylindrical mold, starting with a partially crosslinked gel disc with a 15mm diameter and approximately 0.5 mm thick. Pre-wetted templates, comprised of a support frame, electrospun fibers, and PCL plugs, were positioned on top of this layer and an additional hydrogel was added and crosslinked. After dissolving the template in acetone, the hydrogels were rehydrated and found to swell by 19 ± 5 %. Bright field microscopy of the PEGDA-CO constructs revealed microchannel structures between the two mini-wells (Figure 2C); the nanofibers of the PEGDA-CF device prevented direct evidence of microchannel formation (Figure 3B). To confirm connectivity between mini-wells, a suspension of 0.2-µm-diameter fluorescent microspheres was applied to one mini-well of both the PEGDA-CO and PEGDA-CF devices. The fluorescent microspheres diffused across the microchannel network of both devices within minutes, revealing clearly delineated channels connecting the two wells within an otherwise impermeable hydrogel (Figure 2D, 3C). Confocal microscopy was used to image fluorescent microspheres within PEGDA-CO devices (Figure S5) and confirmed the presence of rounded microchannels with an average diameter of ~6.6 ± 0.78 μm (*n*=8), statistically similar to the original fiber template size. SEM analysis of a cross-section of the PEGDA-CF hydrogel revealed the presence of microchannels with oriented nanofibers around the microchannel periphery (Figure 3E,F). In contrast, PEGDA-FO constructs did not exhibit diffusion of the fluorescent microspheres (data not shown), confirming the formation of a monolithic hydrogel around the encapsulated fibrous scaffold; this indirectly corroborates theoretical calculations of a hydrogel mesh size of approximately 8 nm.^[32]^

To create collagen hydrogel devices, gelation dynamics were measured via parallel plate rheology and collagen crosslinking was observed to be almost complete within 10 min after heating to 37 °C. Therefore, the first layer of collagen was heated for only 5 min to create a partially gelled layer, after which the sacrificial template and/or structural nanofibres were placed on top and a second gel layer was immediately added. After complete gelation, the sacrificial template was fully enscapsulated within a monolithic hydrogel approximately 2 mm thick. Placing the hydrogel device in acetone dissolved the template, forming mini-wells and microchannels while leaving the nanofibers and the hydrogel intact. The acetone treatment also densified the gel, with the final hydrated devices having an approximate thickness of 1 mm. The fragility and opacity of the collagen gel prevented direct confirmation of microchannel patterning via SEM or optical microscopy. However, microchannel formation was indirectly confirmed by the observation of directed DRG outgrowth.

Rat sensory nerve Dorsal Root Ganglia (DRGs) were explanted and placed in all three types of devices (CO, FO, CF). As expected, all collagen-based devices were able to support cell adhesion while only the FO and CF variants of the PEGDA-based devices sustained cell growth, attributed to the presence of nanofibers. Assessment of neurite outgrowth by immunocytochemical (ICC) staining of *β*-Tubullin III (Figure 4B–E) revealed that all nanofiber-containing devices experienced neurite outgrowth in the fiber direction, regardless of hydrogel formulation. However, neurite growth in PEGDA-FO and collagen-FO constructs was confined to the mini-well, with no infiltration of the hydrogel evident (Figure 5B–C).

**Figure 4.**
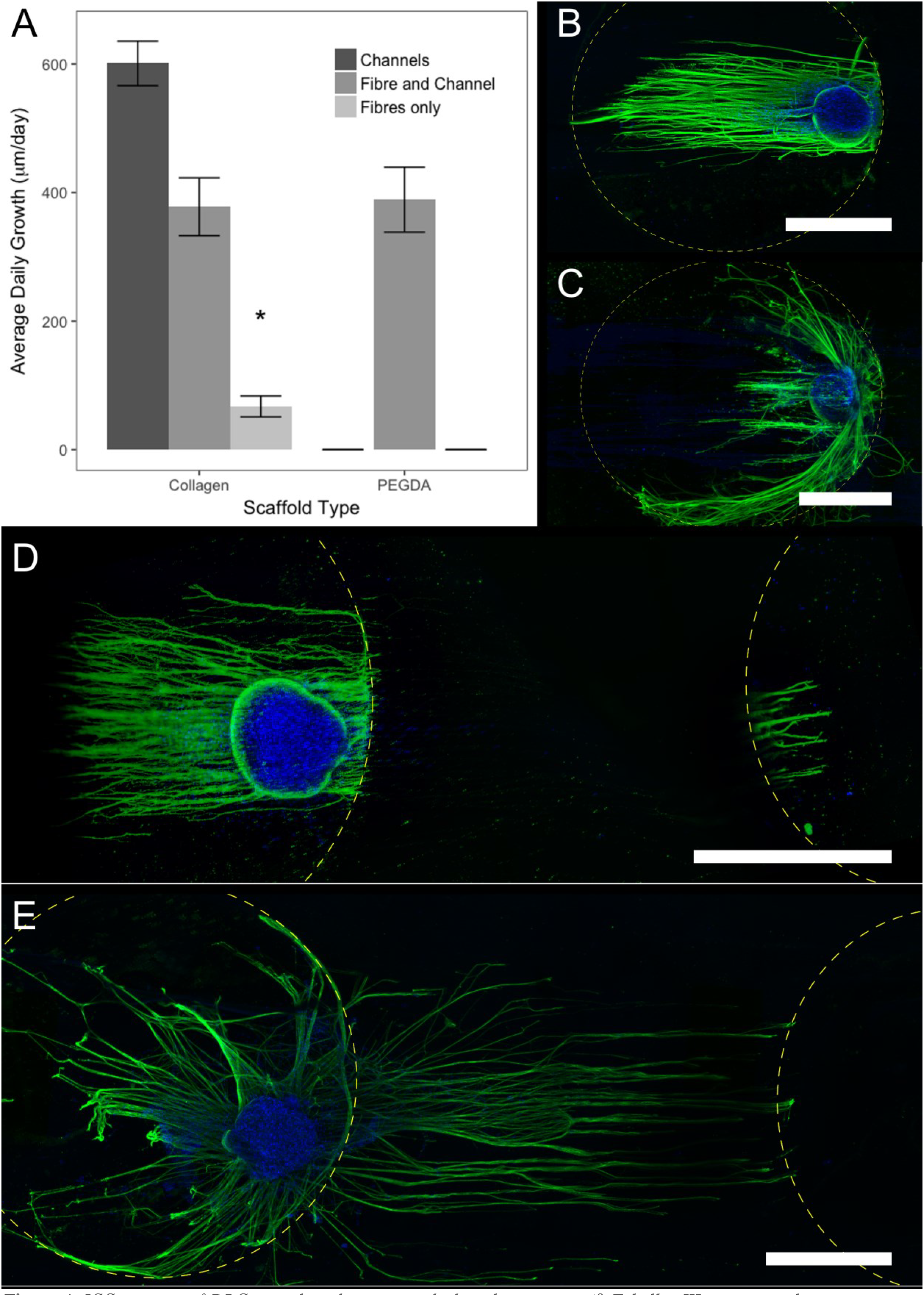
ICC staining of DRG growth within various hydrogel constructs (β-Tubullin III, green; nuclear counterstain (DAPI), blue). Quantification of growth in μm/day show that the presence of channels promotes significantly more growth compared to no channels, provided cells can sufficiently adhere (A). Neurite growth in FO devices was significantly less compared to CF devices, while PEGDA-CO devices could not sustain cell adhesion. No significant differences were found between PEGDA-CF, collagen-CF, and collagen-CO devices; errorbars show SEM (n≥3). Limited neurite growth was observed within both PEGDA-FO devices (B) and collagen-FO devices (C). When PEGDA devices were created with CF templates (microchannels and nanofibers), extensive neurite infiltration was observed (D). ICC staining in these devices exhibited a decreasing gradient with increasing distance from either mini-well, attributed to the limited diffusion of relative large antibodies (20 to 40 nm in diameter) through the PEGDA gel with a calculated mesh size of 8 nm.^[32]^ Continuous neurite growth was confirmed via IB4 lectin (114 kD) and fluorophore-conjugated streptavidin (50 kD), each smaller than 5 nm (Figure S7).^[62]^ Neurite growth was also observed in both collagen-CF and collagen-CO devices (E). Scale bars, 1 mm.

In contrast, all CF devices readily supported neurite growth into the monolithic hydrogel, traversing the microchannel network and entering the adjacent mini-well. Neurite growth through PEGDA-CF devices was indirectly observe through ICC (Figure 4D) and confirmed by an IB4 lectin staining (Figure S7); to ensure the effectiveness of the lectin staining, NGF was substituted with GDNF to promote the growth of IB4^+^ sensory neurons.^[34]^ Both the collagen-CF and PEGDA-CF devices showed similar growth rates, with respective average rates of 377 ± 179 and 389 ± 252 µm/day for the NGF condition (Figure 4A) and 541 ± 320 and 532 ± 304 µm/day for the GDNF condition (Figure S8). The only CO constructs able to support neurite growth were collagen-based, with a growth rate of 600 ± 103 µm/day (Figure 4E); no significant differences were found in growth rates between the CF devices and collagen-CO devices. Without nanofibers present in the collagen-CO devices, the microfibers bundled together during the wetting process and formed larger channels, resulting in less delineated and less organized neurite growth (Figure 5E). Meanwhile, the addition of nanofibers appears to preserve the microchannel structure, further mimicking the role of native collagen fibrils to mechanically stabilize the PNS ECM microarchitecture.^[35]^

## Discussion

The creation of 3D *in vitro* platforms that exhibit representative microenvironments represents a nexus between classical 2D *in vitro* studies and animal models, providing an invaluable tool to study cell behavior in a more relevant context. 2D *in vitro* studies remain valuable to study neurite growth and have been crucial in revealing how the underlying substrate affects cell adhesion and how growth cones are subsequently guided via topographical cues^[36,37]^ and diffusible cues.^[38]^ However, changes in cell adhesion, cell mobility, and cell shape have been described between 2D and 3D environments,^[43,44]^ with differences specifically noted for growth cones morphology and nerve growth.^[45]^ This has motivated the development of 3D *in vitro* culture system, many of which are already in use in neuroscience studies. However, attempts to emulate the native ECM have comparatively large microchannel structures,^[11,50]^ lack nanofibrous topographical guidance,^[27,51,52]^ or employ specialized techniques that limit widespread adoption.^[53]^ Other techniques offer promising approaches to produce microchannel networks within a hydrogel, including Omnidirectional Printing and Melt Electrospinning Writing (MEW), but have yet to be applied to neural applications.^[54,55]^ In contrast with previously reports,^[26,27]^ the sacrificial template strategy outlined here creates microchannels delineated with aligned nanofibers to thoroughly emulate the PNS structure/function relationship of native ECM endoneural tubes lined with oriented collagen fibrils.

A comparison of reports provides insight into the impact of different *in vitro* culture environments on nerve growth, though care must be taken to account for culture conditions and growth metrics when assessing neurite growth across different studies. 2D studies were the first to examine neural growth on softer surfaces, using 2D hydrogel films to identify an optimal substrate stiffness above 1 kPa^[39,40]^ which can achieve growth rates of 100 to 130 µm/day to match the 100 to 140 µm/day observed on glass or TCP surfaces.^[41,42]^ In contrast to 2D substrates, neurite growth within featureless 3D collagen hydrogels exhibit faster growth rates ranging from 200 to 260 µm/day. ^[20,33,46]^ Further improvements are reported for neurites grown on oriented nanofiber substrates, with this partial mimic of the ECM structure resulting in growth ranging from 280 to 650 µm/day;^[18,47]^ although these substrates may be considered closer to 2D environments, studies suggest that incorporating nanotopographical cues can also impact 3D scaffold performance. Oriented synthetic nanofibers embedded within collagen have produced neurite growth of 290 µm/day,^[46]^ while directly aligning the collagen fibrils intrinsic to the hydrogel resulted in highly oriented neurite extension and a significant increase of 370 µm/day.^[20]^ This is in line with evidence that fibril orientation can have a greater impact on cell adhesions than hydrogel stiffness.^[48]^

However, gel stiffness still plays an important role within a 3D context, as observed by a 0.5 kPa hydrogel achieving neurite growth of 330 µm/day while a 2.1 kPa gel experiencing markedly reducing growth to 85 µm/day.^[2]^ Running contrary to observations of neurites on 2D hydrogels, the reduction in growth can be attributed to higher density gels requiring more active remodeling by cells.^[2,21,49]^ This has motivated the incorporation of microchannels within gels to create a more permissive environment. While the previously mentioned nerve guide with 3D microchannels experienced *in vitro* neurite growth rate of 145 µm/day, fibrin hydrogels with micron-scale channels exhibited growth rate of approximately 500 um/day.^[27]^ Overall, these reported findings highlight the impact of different elements of ECM architecture on neurite growth.

The hybrid scaffold reported here represents a significant improvement over previous 3D culturing environments, with an organized nanofibrillar microarchitecture that emulates the *in vivo* ECM structure to produce rapid and directed neurite growth on a time scale approaching that observed *in vivo*. Axons maintained unobstructed growth via the oriented microchannel network at rates of up to 390 µm/day over a 5-day period. In the presence of GDNF, both collagen-CF and PEGDA-CF devices achieved an average growth rate of 500 µm/day, with one PEGDA-CF device exhibiting a remarkable growth rate of 900 µm/day.

Unique to this culturing platform is the ability to effectively decouple fibril orientation, material density, and substrate stiffness within a 3D environment. This makes it possible to deconvolve the impact of the microenvironment on cellular response, towards better understanding and more effective therapies. With the obvious caveat that the nanofiber and hydrogel materials must withstand the template removal process, this device provides the possibility to systematically assemble different synthetic ECM configurations. Only one type of nanofiber was evaluated in the current study. However, electrospun fibers can influence cell behavior through different material compositions, dimensions, alignment and density, thus expanding the possible use of this platform to other tissues.^[29]^ Furthermore, a number of other hydrogel materials are known to be stable in acetone include fibrin,^[27]^ agarose,^[30]^ and pHEMA.^[31]^ With regards to the use of natural hydrogel materials and previous reports of cells remodeling this 3D environment,^[33]^ neurites in the current study were instead observed to follow the microchannel structures. This was possibly because of collagen densification during acetone exposure, thus limiting the ability to remodel the matrix, or axons simply taking the path of least resistance. Overall, the processing steps described resulted in the robust production of an in vitro platform specifically designed for directed neurite growth 3D.

With the emergence of 3R regulations (replacement, reduction, and refinement) for animal use in research, the development of relevant 3D *in vitro* platforms are becoming critical to allow scientists to continue to address important biological question.^[56]^ With peripheral nerve injury (PNI) occurring in approximately 2.8% of all trauma injuries^[57]^ and resulting in reduced patient quality-of-life,^[58]^ 3D *in vitro* platforms that provide an *in vivo*-like environment capable of exploring the finer nuances of neurite growth will become vital to improving nerve repair strategies. Furthermore, the current two-well system connected via well-defined microchannel structures also lends itself to creating 3D neural circuits with more directed organization than previously reported.^[59–61]^ While our application focused on studying neurite growth, this robust and versatile platform can also explore other cell behavior, such as migration or vascularization within such a pre-patterned network.

The current study presents a highly flexible synthetic analog to the native endoneurium, achieving for the first time passive guidance of extensive nerve growth via a combination of corralling microstructure and oriented fibrous nanotopography within a compliant 3D environment. The ability to create a hygroscopic environment that combines aligned fibers with directed microchannel porosity in a controlled manner realizes a scaffold design with a new degree of order and complexity. In addition, we present a modular fabrication strategy able to implement structural elements within both synthetic and natural hydrogels, facilitating the controlled study of cell response to multivariate 3D environments. This platform clearly shows the significance of incorporating the neuromimetic ECM architecture, which stimulates similar nerve growth in both PEGDA and collagen despite inherent differences between these natural and synthetic materials. The resulting 3D culturing platform is compatible with standard *in vitro* culturing practices and is amenable to microscopy analysis, providing new opportunities to study mechanisms of nerve growth and to optimize tissue repair strategies.

## Acknowledgments

The authors would like to thank H. Nyugen for critical revision of the manuscript and to acknowledge the funding support of the Natural Sciences and Engineering Council of Canada (NSERC), the European Union FP7-NMP project MERIDIAN under contract number 280778, and the Province of Limburg, the Netherlands.

