## Supplementary Data for "Meso-scale multi-material fabrication of a Synthetic ECM Mimic for In vivo-like Peripheral Nerve Regeneration"

### **Supporting Information:** Biomimetic Nanofiber-Hydrogel Hybrid Construct

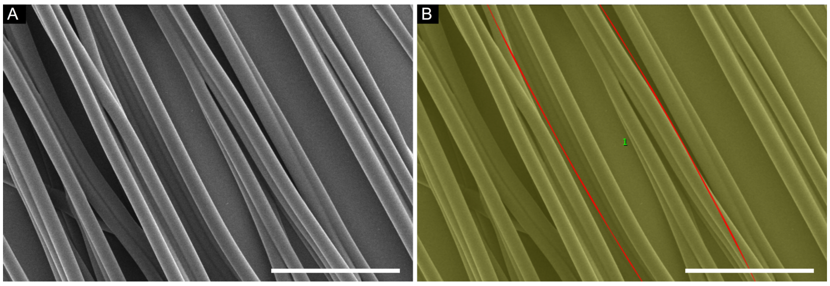


***Figure S1.*** *An example image of assessment of fiber alignment. The original image of fibers produced with 50% w/v PLA at a flow rate of 5 ml/hr (A) is processed with the OrientationJ plugin (B), available for ImageJ. This assessed the degree of alignment of features within the image, producing a value of coherence between 0 (random) and 1 (aligned). (Scalebar: 50 µm)*

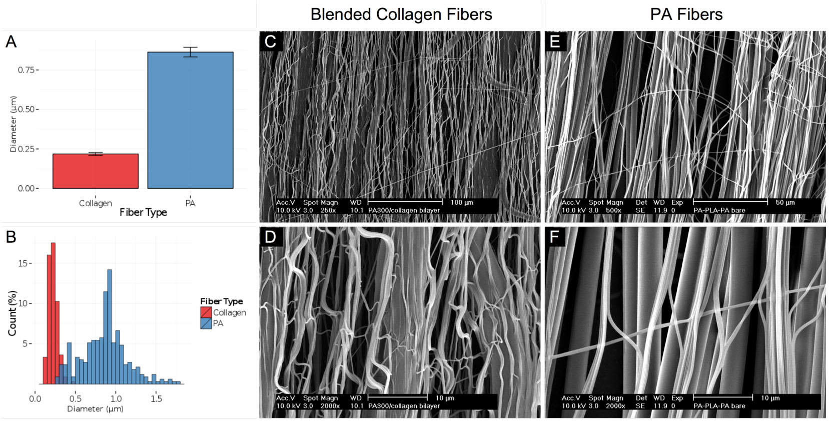


***Figure S2***. *The diameters of both types of insoluble fiber diameter are shown as average values (A) and distributions (B). Pure PA fibers had an average diameter of 0.863±0.281 µm, while blended PA-collagen fibers were a 0.218±0.057 µm. These fibers were used to create either triple layer fiber constructs of either Collagen/PLA/Collagen (C, D) or PA/PLA/PA (E, F).*

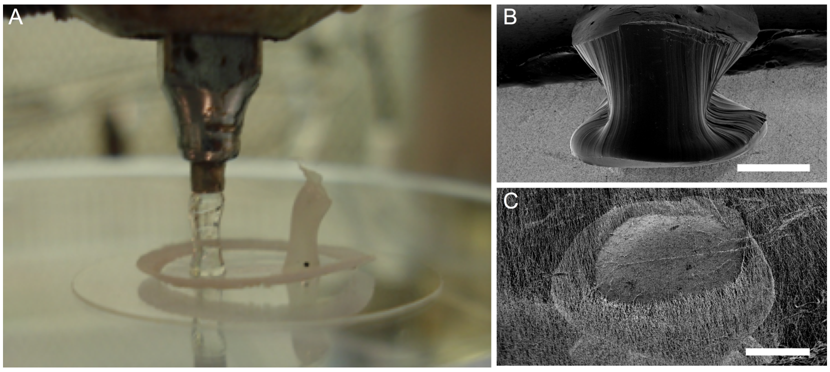


***Figure S3.*** *PCL plug integration with the fiber network. The PCL plug was deposited on the fiber network via a bioplotting device (A). The molten plug was able to adhere to the fiber mesh surface (B), penetrating through the fiber mesh (C). (Scalebar: 1 mm)*

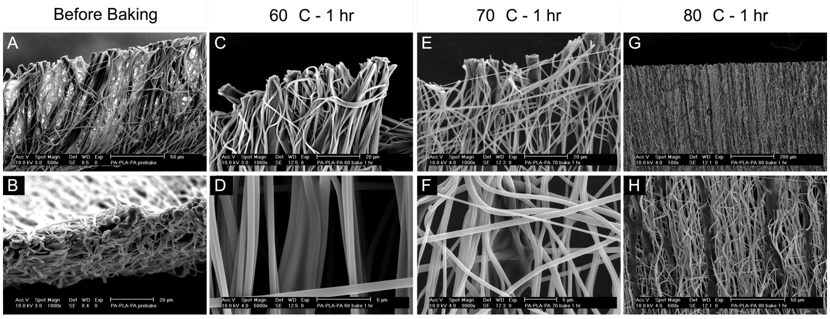


***Figure S4.*** *Optimization of the merging of PA and PLA fibers by the baking. Before the application of heat, the two fiber types are distinct (A, B). After the application of 60 °C for one hour (C, D), slight softening of the PLA fibers can be observed but the degree of merging is limted. Baking for either 70 °C (E, F) or 80 °C (G, H) for one hour increases the degree of merging between the fiber types, however the PLA fibers lose their rounded morphology.*

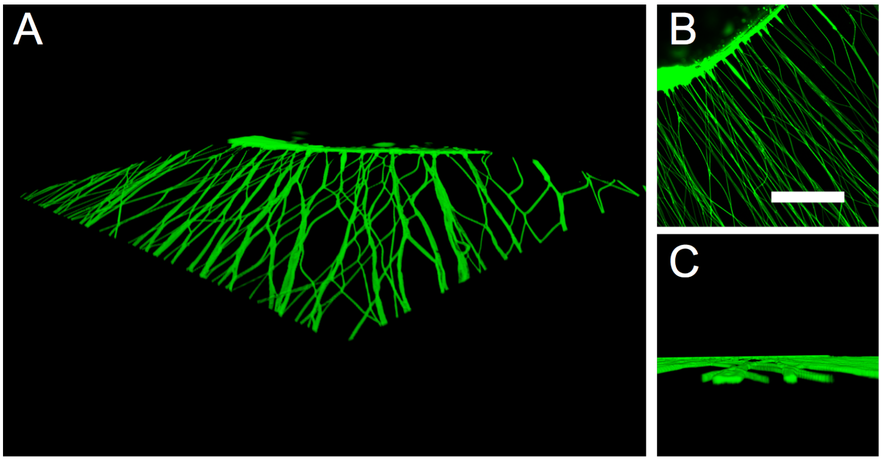


***Figure S5****. A 3D rendering (A) of suspended fluorescent nanospheres as they flow from the seeding well to fill microchannels formed within a PEGDA-CO hydrogel, acquired via confocal microscopy (B). The morphology of the microchannels (C) retains the shape of the original template fibers. Scale bar: B, 400 µm.*

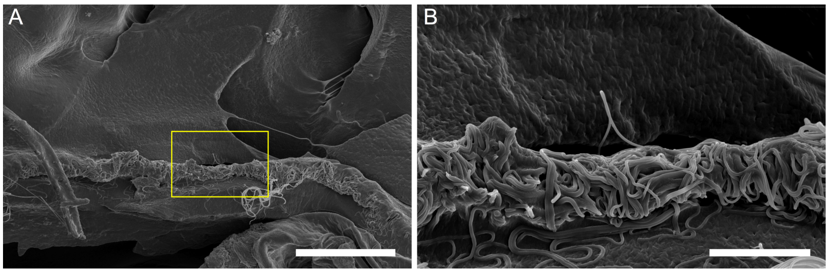


***Figure S6***. *Cross sectional view of a hydrogel device with the typical density of insoluble fibers (A). At this high density, the presence of microchannel structures is obscured by the insoluble fibers (B). (Scalebar: A, 100 µm; B, 20 µm)*

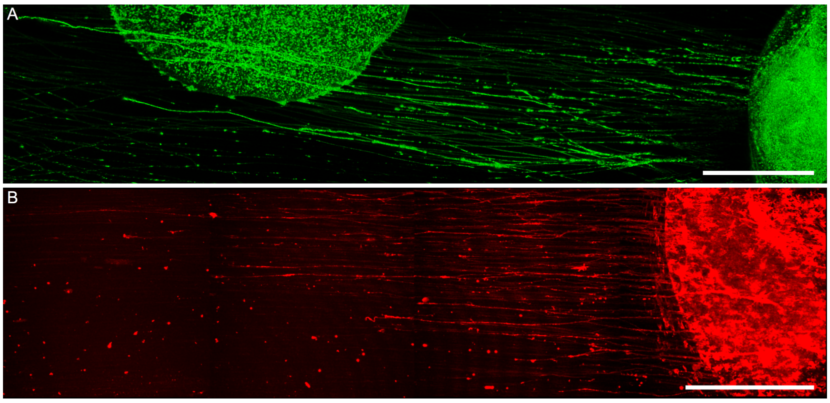


***Figure S7.*** *Confirmation of neurite growth throughout gel using lectin staining for IB4+ of neurites grown in NGF. (A) Neurite outgrowth of DRGs explanted from post-natal Day 8 rat pups, grown in NGF over an 8 day period. (B) neurite outgrowth of DRGs explanted from post-natal Day 2 rat pups, grown in NGF over a 5 day period. (Scalebar: 500 µm)*

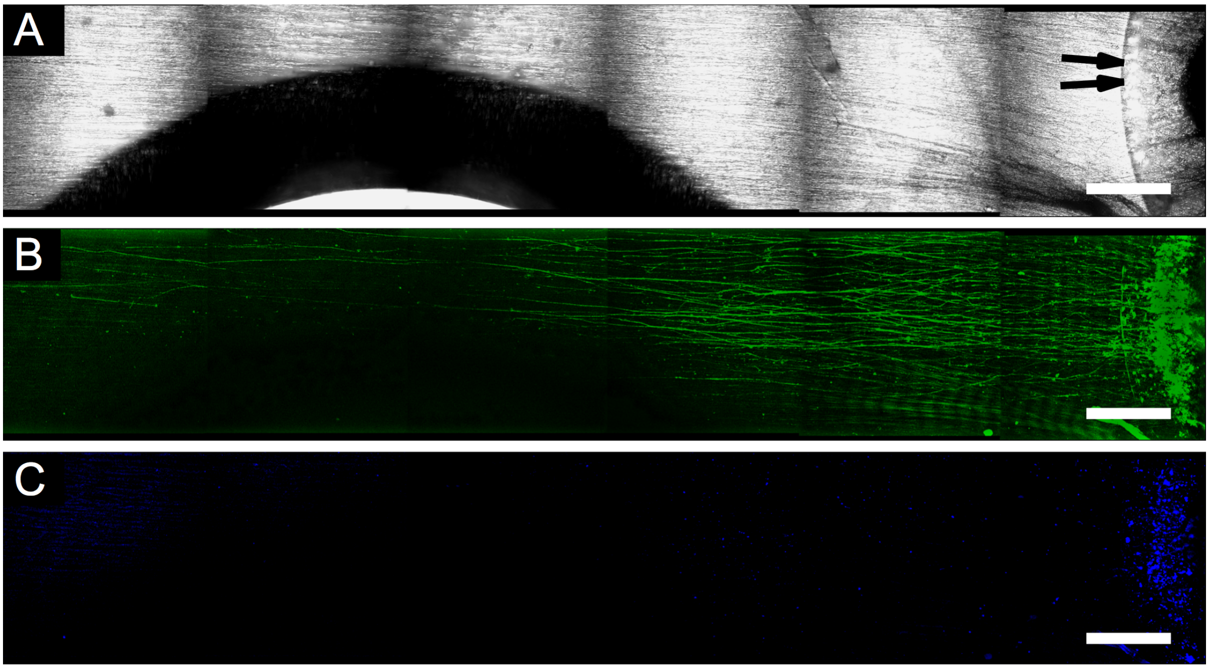


***Figure S8.*** *Confirmation of neurite growth throughout gel using lectin staining for IB4+ of neurites grown in GDNF. (A) A brightfield image of a PEGDA-CF device, showing the perimeter of the mini-well (arrows). Also visible is an air bubble trapped under the PEGDA. (B) Neurite outgrowth of DRGs explanted from post-natal Day 8 rat pups, grown in NGF over an 8 day period, fixed, and stained for IB4+ neurites. Neurite growth over a distance of 7 mm was observed, approaching the 1 mm/day growth rate observed in vivo. (C) DAPI staining showing the confinement of cell to the mini-well region. (Scalebar: 500 µm)*

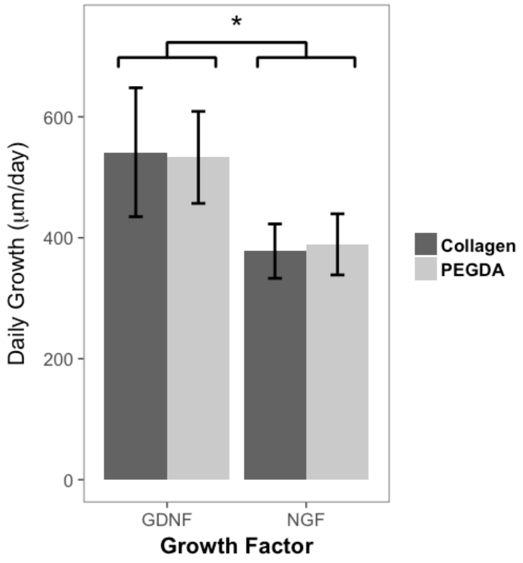


**Figure S8.** Comparison of neurite growth rates in CF constructs. Under a given growth factor, no difference is observed between collagen and PEGDA CF constructs. GDNF is found to promote significantly faster neurite growth compared to NGF.
